# BayesAge: A Maximum Likelihood Algorithm To Predict Epigenetic Age

**DOI:** 10.1101/2023.12.03.569788

**Authors:** Lajoyce Mboning, Liudmilla Rubbi, Michael Thompson, Louis-S. Bouchard, Matteo Pellegrini

## Abstract

DNA methylation is a reaction that results in the formation of 5-methylcytosine when a methyl group is added to the cytosine’s C5 position. As organisms age, DNA methylation patterns change in a reproducible fashion. This phenomenon has established DNA methylation as a valuable biomarker in aging studies. Epigenetic clocks based on weighted combinations of methylation sites have been developed to accurately predict the age of an individual from their methylome. However, many epigenetic clocks, particularly those that utilize penalized regression, model the changes in methylation linearly with age. Moreover, these models, which use methylation levels as features, are not robust to missing data and do not account the count-based nature of bisulfite sequence data. Additionally, the models are generally not interpretable. To overcome these challenges, we present BayesAge, an extension of the previously developed scAge approach that was developed for the analysis of single cell DNA methylation datasets. BayesAge utilizes maximum likelihood estimation (MLE) to infer ages, models count data using binomial distributions, and uses LOWESS smoothing to capture the non-linear dynamics between methylation and age. Our approach is designed for use with bulk bisulfite sequencing datasets. BayesAge outperforms scAge in several respects. Specifically, BayesAge’s age residuals are not age associated, thus providing a less biased representation of epigenetic age variation across populations. Moreover, BayesAge enables the estimation of error bounds on age inference and, when run on down-sampled data, its coefficient of determination between predicted and actual ages surpasses both scAge and penalized regression.

## 1 INTRODUCTION

While the sequence of a cell’s DNA largely remains invariant during its lifespan, its epigenome changes significantly with age. One of the components of the epigenome that shows the most reproducible changes with age is DNA methylation. DNA methylation involves the covalent modification of DNA and is catalyzed by a family of DNA methyltransferases (DNMTs) that transfer a methyl group from S-adenosyl methionine (SAM) to the fifth carbon of a cytosine residue to form 5-methylCytosine (5mC).Moore et al. (2012) In mammals, most of the methylated cytosines occur in the CpG context, and there are approximately 30M 5’—C—phosphate—G—3’ (CpG) dinucleotides in the human genome.

Several methods have been developed to predict age based on methylation levels.Johnson et al. (2012) These approaches are often referred to as epigenetic clocks.Kabacik et al. (2022)The first epigenetic clocks were constructed using penalized regressionGreenwood et al. (2020), with age as the response and methylation levels as the features. The use of penalties for model coefficients led to sparse models out of the thousands of sites that were measured (typically using DNA methylation microarrays), only a few hundred had non-zero weights in the models.Hannum et al. (2013); Horvath (2013) More recently these approaches have been extended using neural networks, which can generate slightly more accurate predictions than penalized regression models.Galkin et al. (2021)However, these approaches model DNA methylation changes linearly with age and therefore fail to consider non-linear methylation trends with age. This is significant, as it is widely understood that methylation changes are rapid early in life and slow down with age.Snir et al. (2019)

To address this limitation, Farrell and colleagues introduced a model named the Epigenetic Pacemaker (EPM).Farrell and Pellegrini (2020) In this model, methylation across a subset of CpGs is described as a non-linear function of an epigenetic state, rather than actual age. Importantly, this epigenetic state can assume non-linear relationships between the with age. For a given set of *i* methylation sites and *j* individuals, the methylation level at a single site can be expressed as 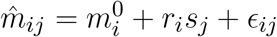, where 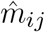 represents the observed methylation value, 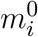 denotes the initial methylation level, *r*_*i*_ is the rate of change, *s*_*j*_ signifies the epigenetic state, and *ϵ*_*ij*_ is a normally distributed error term.

Given an input matrix 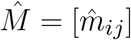, the objective of the Epigenetic Pacemaker (EPM) is to ascertain the optimal values for 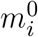, *r*_*i*_, and *s*_*j*_ that minimize discrepancies between predicted and actual methylation values across specific methylation sites. As we’ve previously demonstrated, in certain datasets, the epigenetic state evolves in correlation with the logarithm of time.Snir et al. (2019) This implies rapid methylome alterations early in development, which then decelerate with the organism’s aging. Although the EPM adeptly models some nonlinear correlations between methylation and age, it calibrates the epigenetic age in relation to chronological age to optimize alignment across all sites. However, it overlooks the potential nonlinear associations that different sites may exhibit with age.

Here we propose a new approach to overcome the limitations of existing methods. Our method considers count information rather than methylation fractions, estimates non-linear trends of methylation sites with age, and uses maximum likelihood estimation which is robust to missing data. Existing methods for methylome aging modeling often struggle with low coverage or incomplete data. For instance, linear models for epigenetic age, built on weighted sums of methylation values, inadequately handle missing data—typically by assigning it a value of zero—thereby skewing age predictions. Addressing this deficit, maximum likelihood methods have emerged to generate age estimates from sparse data sets. One existing method, “scAge” (single cell age),Trapp and Gladyshev (2021) is designed to analyze methylation in single cells. scAge harnesses a maximum likelihood strategy to ascertain the most likely age of a subject based on low count data. However, this method presumes a linear relationship in methylation changes over time and employs a heuristic for determining methylation value probabilities from the observed data. We therefore expand upon scAge’s foundational principles to develop a read count based framework for modeling non-linear methylation changes with age that is resilient against missing data.

## 2 MATERIALS AND METHODS

### 2.1 Data acquisition and analysis

To test our approach for estimating age from methylation data we collected targeted bisulfite sequencing data from either buccal swabs or blood of 458 subjects. DNA was extracted from the buccal swabs and blood using standard protocols. Buccal swabs were incubated overnight at 50°C before DNA extraction. We applied targeted bisulphite sequencing (TBS-seq) to characterize the methylomes of the samples. The protocol is described in detail in a methods paper by Morselli et al.Morselli (2021)Briefly, 500 ng of extracted DNA were used for TBS-seq library preparation. Fragmented DNA was subject to end repair, dA-tailing and adapter ligation using the NEBNext Ultra II Library prep kit using custom pre-methylated adapters (IDT). Pools of 16 purified libraries were hybridized to the biotinylated probes according to the manufacturer’s protocol. Captured DNA was treated with bisulphite prior to PCR amplification using KAPA HiFi Uracil+(Roche) with the following conditions: 2 min at 98°C; 14 cycles of (98°C for 20 sec; 60°C for 30 sec; 72°C for 30 sec); 72°C for 5 minutes; hold at 4°C. Library QC was performed using the High-Sensitivity D1000 Assay on a 2200 Agilent TapeStation. Pools of 96 libraries were sequenced on a NovaSeq6000 (S1 lane) as paired-end 150 bases. The probes used in the capture were designed to capture approximately 3000 regions that contained CpG sites used in previously published epigenetic clocks. Greenwood et al. (2020); Levine (2018); Lu (2019)

Demultiplexed Fastq files were subject to adapter removal using cutadapt (v2.10)Martin (2011) and aligned to the GRCh38 genome using BSBolt Align (v1.3.0)Colin (2021). PCR duplicates were removed using samtools markdup function (samtools version 1.9)Li (2009). The BSBolt methylation calling function was employed to produce the CGmap files for each subject, utilizing the sorted and indexed bam files. The reference genome used during the methylation calling phase was Genome assembly GRCh38. The BSBolt matrix aggregation function was used to create the methylation matrix dataset, which subsequently informed the training of BayesAge, scAge, and the LASSO model. Methylation values were measured across 46,518 CpG sites.

CGmap files were converted to Bismark format for scAge testing. Additionally, BayesAge incorporates a function to convert CGmap files into a format compatible with its prediction function.

The LASSO model was implemented using the RepeatedKFold package from `sklearn` in Python. The parameters used to train the lasso model had an alpha value equal to 0.02 and max iterations of 10,000. The prediction parameter of the lasso model had a cross validation of 10. Due to extreme downsampling to 100,000 CpG sites, the methylation calling using BSBolt would return empty matrices. To circumvent this issue, a methylation matrix with very low values of coverage and percent of samples was used. Since the Lasso model requires finite input values, any NaN entries were imputed as zero methylation. While not ideal, this allowed the model to be trained and make age predictions on the available data.

### 2.2 The BayesAge framework

The BayesAge framework consists of two phases: traning and prediction. In the training step, we use LOWESS (locally weighted scatterplot smoothing) to fit the trend between individual methylation levels and age. We use LOWESS smoothing so that we do not need to make any a priori assumptions about the functional form of the association between methlyation and age. The *τ* parameter determines the smoothness of the LOWESS fit. The LOWESS function used was from the `statsmodels.api`. After computing the fit for each site we also calculate the correlation between methylation levels and age using the Spearman rank correlation, which is robust to non-linear trends. We select the top sites to include in the prediction phase using the absolute value of the correlation. At the end of this process, the trained model consists of *N* sites and their methylation levels across ages, from 1 to 100 in increments of one year, based on a predetermined *τ* parameter of the LOWESS fit.

In the Prediction step, this reference matrix is intersected with the CpG sites measured in a specific sample. A count matrix of these CpG sites is constructed that reports the number of cytosines and thymines. For the chosen age-associated CpG sites, it’s posited that the chance of detecting the observed cytosine and thymine counts given the intended methylation level for a specific age based on the trained model, follows a binomial distribution. To compute the probability of observing the counts measured across all sites that are found following the intersection with the training matrix, we compute the product of these probabilities. To prevent underflow errors during computation, a logarithmic sum replaces the product of individual CpG probabilities, which results in a singular probability value for each age.

Utilizing these pre-identified, ranked age-associated CpG sites, the framework calculates the likelihood of observing each age in a single subject, spanning an age spectrum of 0 to 100 years, at an interval of 1 year. Consequently, for each subject we compute an age-likelihood distribution, with the maximum likelihood age interpreted as the epigenetic age for subject *X*. Here, *Pr*_*CpG*_ represents the methylation probability for a distinct CpG at a specific age, aggregated from 1 CpG to *N* total CpGs. The associated probability for a unique CpG site state is mathematically detailed as follows:

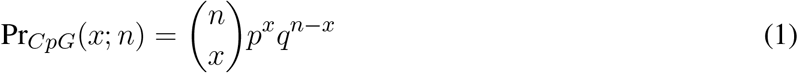

where:

*n*: Reads of all cytosines.

*x*: Reads of methylated cytosines.

*p*: The predicted average methylation probability.

*q* =1 − *p*: The probability of thymine counts.

#### 2.2.1 Data Simulation and Average

The data simulation was executed by creating 100 synthetic samples for each real sample for a total 45800 samples. For each synthetic sample, the counts at each site were simulated using the `scipy.stats.binom` probability distribution function. Futhermore, BayesAge prediction function was used to estimate the age of each of the 100 synthetic samples. The 100 synthetic samples of each real sample was used to calculate the lower interquartile range (IQR) and upper IQR limit bounds. Finally, the average age of the error limit bounds reported were calculated using:

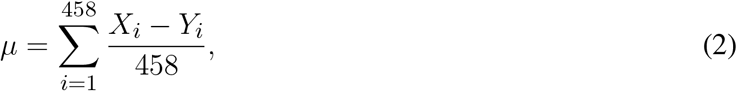

where *X* and *Y* represent the upper IQR limit and lower IQR limit of sample *i* respectively.

## 3 RESULTS

### 3.1 BayesAge framework

We set out to develop a framework to estimate the age of an individual from bisulfite sequence data. Bisulfite conversion converts unmethylated cystosines to thymines, while leaving methylated cytosines unconverted. After bisulfite converted DNA is sequenced, the reads are aligned to genome and the methylation state of any cytosine in the genome is measured by counting the number of cytosines and thymines that align to that position. Typically the methylation level is estimated by computing the ratio of cytosines to cytosines plus thymines. However, since sequencing data is inherently count based, it is important to consider not only the methylation level, but also the total coverage, as the confidence of the methylation estimate increases with increased coverage.

To track changes in DNA methylation with age we collected DNA methylation data from over 400 individuals. To identify CpG sites whose methylation changed with age in a tissue independent manner, we collected our sample from both blood, saliva and buccal swabs. These tissues have heterogeneous mixtures of both hematopoietic as well as epithelial cells, and represent typical cells that are found in many tissues. As whole genome sequencing is resource intensive, we used a targeted approach to only sequence a few thousand loci across the genome. This allowed us to obtain about 100x coverage of 46517 targeted regions. We made sure to include among these loci regions that had been previously shown to have age associated DNA methylation changes. We include samples that covered a broad range of ages, from neonates to 92 years old.

Our BayesAge framework has two steps: in the first we train a model and in the second we predict the age of a sample. In the training phase we first select CpG sites in the genome that have age associated methylation values. Previous work has shown that many CpG sites in the genome have DNA methyation levels that increase with age, but that the association between methylation and age is not necessarily linear. In fact many sites show non-linear changes of methylation with age that are well approximated by exponential functions, with the rate of change decreasing with age. To identify the most significantly age-associated sites, we used the Spearman rank correlation, which is a non-parametric method that does not assume linearity. The 16 sites in the genome with the highest Spearman correlation values are shown in Figure 2.

**Figure 1.**
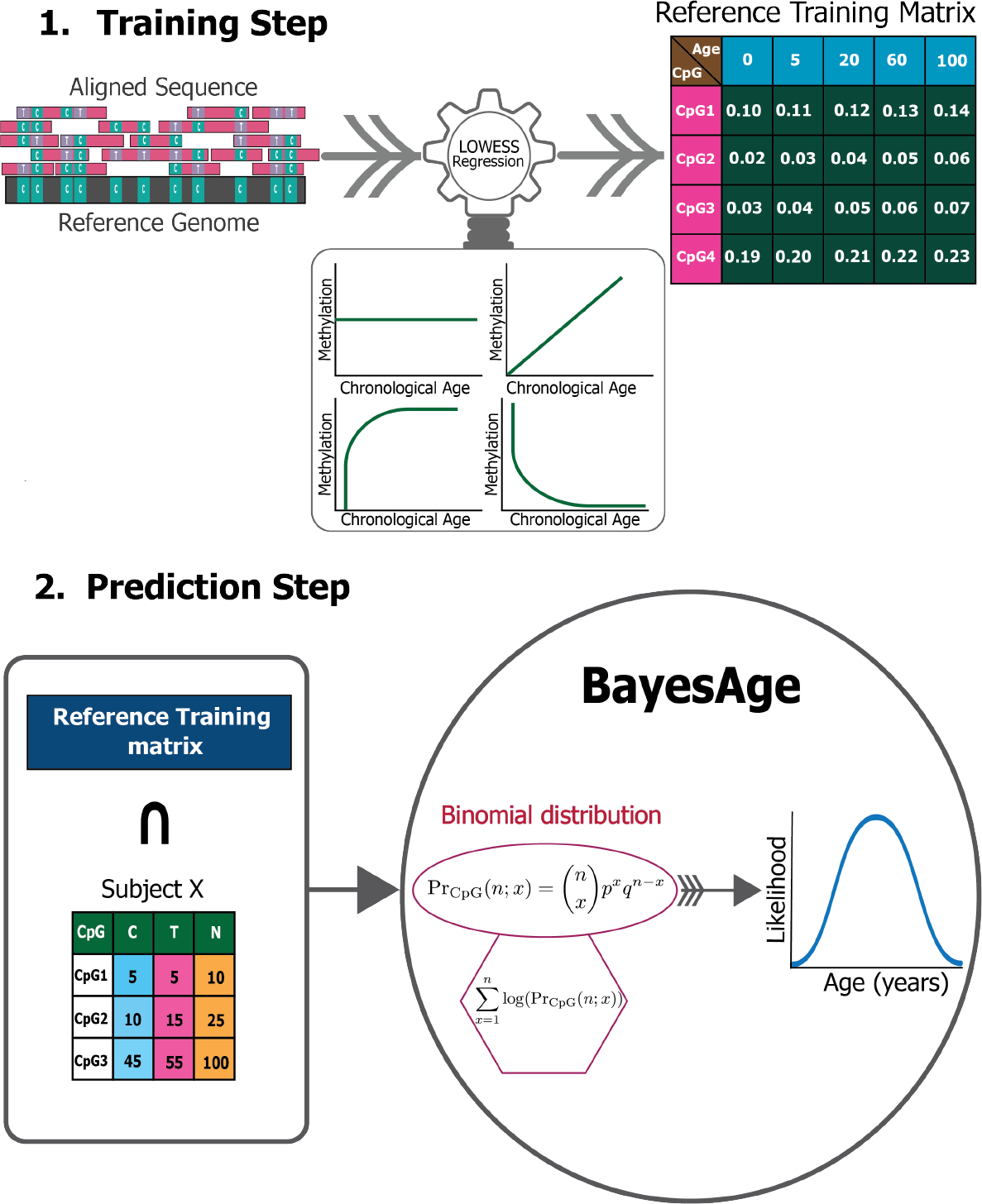
BayesAge framework

**Figure 2.**
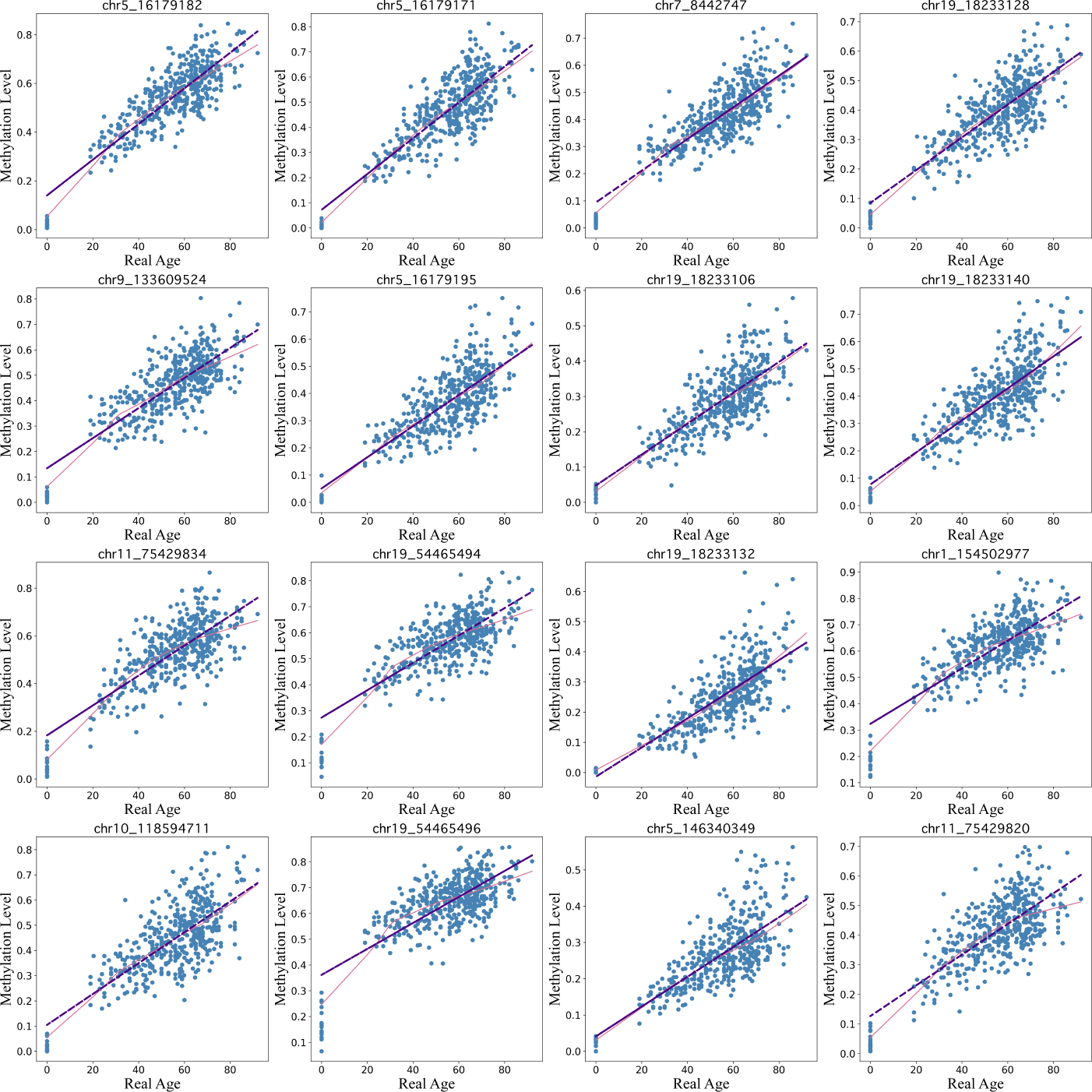
Top 16 CpG sites fitted using lowess regression (pink line) with a tau parameter value of 0.7 vs linear regression (purple line).

We find that for many of these sites the association of methylation with age is non-linear, with rates of change that decrease with age. BayesAge, recognizing these non-linear trends, uses LOWESS (locally weighted scatterplot smoothing) regression to model the trend lines. This method utilizes locally weighted linear regression to estimate a smoothed line to the data. The extent of this smoothing is governed by the tau parameter, which determines the size of the local neighborhood window for each local linear regression fit. For our dataset, a tau value of 0.7 was chosen, allowing the fit to adaptively capture the nonlinear methylation variations across the age spectrum without succumbing to overfitting, as illustrated in Fig. 2. The trend lines of these fits represent our aging model, or the expected methylation with age at each of these sites.

Our prediction step allows us to estimate the age of a sample using the training model. Our approach to estimating age from the methylation data of a single sample builds on the count based nature of bisulfite sequencing data. We propose a Bayesian framework for estimating the most likely age of an individual by computing the probability of the observed counts of cytosines and thymines for any given age, and selecting the age that maximizes this probability. This maximum likelihood approach was first proposed in the scAge method, which was developed to estimate the age of a samples from single cell methylation data. In contrast to scAge, our method is designed for bulk DNA methylation data where the coverage of cytosines in the genome is generally high, and the methylation levels can assume values between zero and one.

To estimate the probability of the observed counts based on the expected methylation levels of a single site at a specific age we use a binomial distribution. The rationale for utilizing a binomial distribution to characterize the sites is anchored in the nature of bisulfite sequencing data. Rather than probabilities, bisulfite sequencing yields count data. Each sequenced read at a CpG site represents a Bernoulli trial with two potential outcomes: the observation of a methylated cytosine, the probability of which equals the intrinsic methylation level at that site, or the observation of an unmethylated cytosine, with the probability being the complement of the methylation level. Consequently, the cumulative counts of methylated and unmethylated reads across all sequenced reads at a particular site adhere to a binomial distribution dictated by the methylation level. This modeling approach for CpG sites echoes the discrete, count-centric nature of sequencing data, diverging from the notion of methylation as continuous probabilities.

Figure 1 and Eq. (1) exemplify the probability calculation phase, which evaluates the likelihood of detecting specific cytosine and thymine counts, given the anticipated methylation level for each CpG across varying ages, as derived from the LOWESS regression. In contrast, scAge’s training phase employs a linear regression model, as evident in Eq. (3), to forecast the methylation level for every CpG site across different ages.

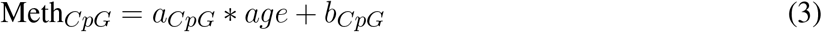

In the final step, in order to estimate the probability of a specific age of a sample we compute the likelihood of our observed counts across multiple sites for any given age by taking the product of the probabilities of each site. In practice this product is computed by summing the logarithms of the probabilities. In the final stage we compute the age that maximizes this probability.

We applied BayesAge to our samples using a model that contains the top 8 sites and used 10-fold cross validation to train and test the model. We find that the *R*^2^ between the predicted age and the actual age is 0.78, and that the mean absolute error of the age estimate is 7 years, indicating a relatively strong correlation and accuracy in age prediction. The cross validation method is implemented such that the model is trained on 448 random samples and then tested on the remaining 10 samples. This process is repeated until all the samples are used for prediction.

Along with estimating the most likely age of a sample, BayesAge enables the calculation of error bounds on the estimate. To generate the prediction error bounds for bdAge, we employed data simulation. By running 100 simulations for each sample, we derived the interquartile range (IQR) of age predictions as a measure of uncertainty. Across all the simulations, the average IQR was approximately 12 years, providing an estimate of the typical error margins for Bayesian age predictions in our dataset as seen in Fig. 3.

**Figure 3.**
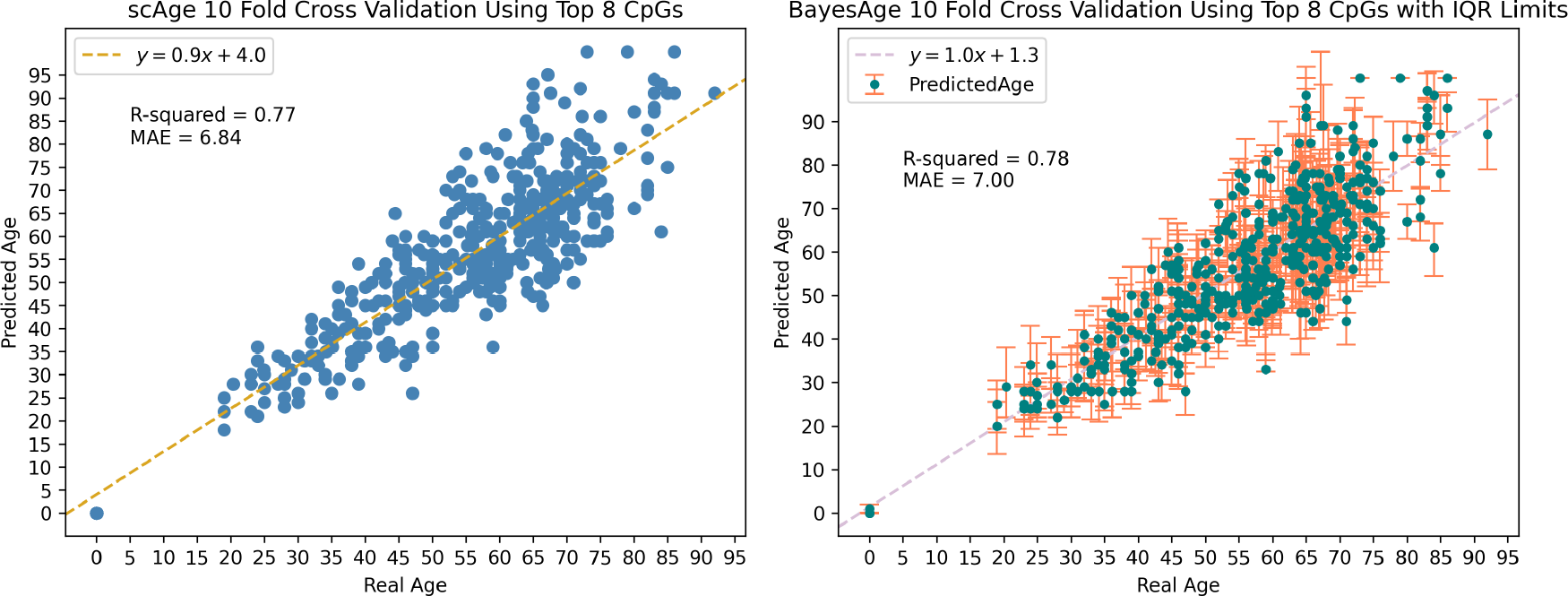
Data simulation was used to generate the IQR limits seen in BayesAge’s plot with a tau value of 0.5.

#### Comparison of BayesAge with scAge and penalized regression models

To evaluate the performance of BayesAge we compared it to scAge and penalized regression models using the same dataset of 458 individuals. As illustrated in Fig. 4, 10-Fold Cross Validation coupled with mean absolute error (MAE) was employed to validate the outcomes of all models. The results show that BayesAge age estimations have a slightly higher coefficient of determination (*r*^2^) compared to scAge. Notably, when limited to the top 8 CpG sites for age prediction, BayesAge outperforms scAge by approximately 1%. However, as we increase the number of CpG sites employed in the prediction, this performance disparity becomes more evident. Specifically, utilizing the top 256 CpG sites, BayesAge achieves an *R*^2^ value of 73%, while scAge lags slightly behind with 70%. It’s worth noting that, across the varying numbers of CpG sites assessed, scAge consistently generated age predictions with slightly lower mean absolute errors relative to BayesAge.

**Figure 4.**
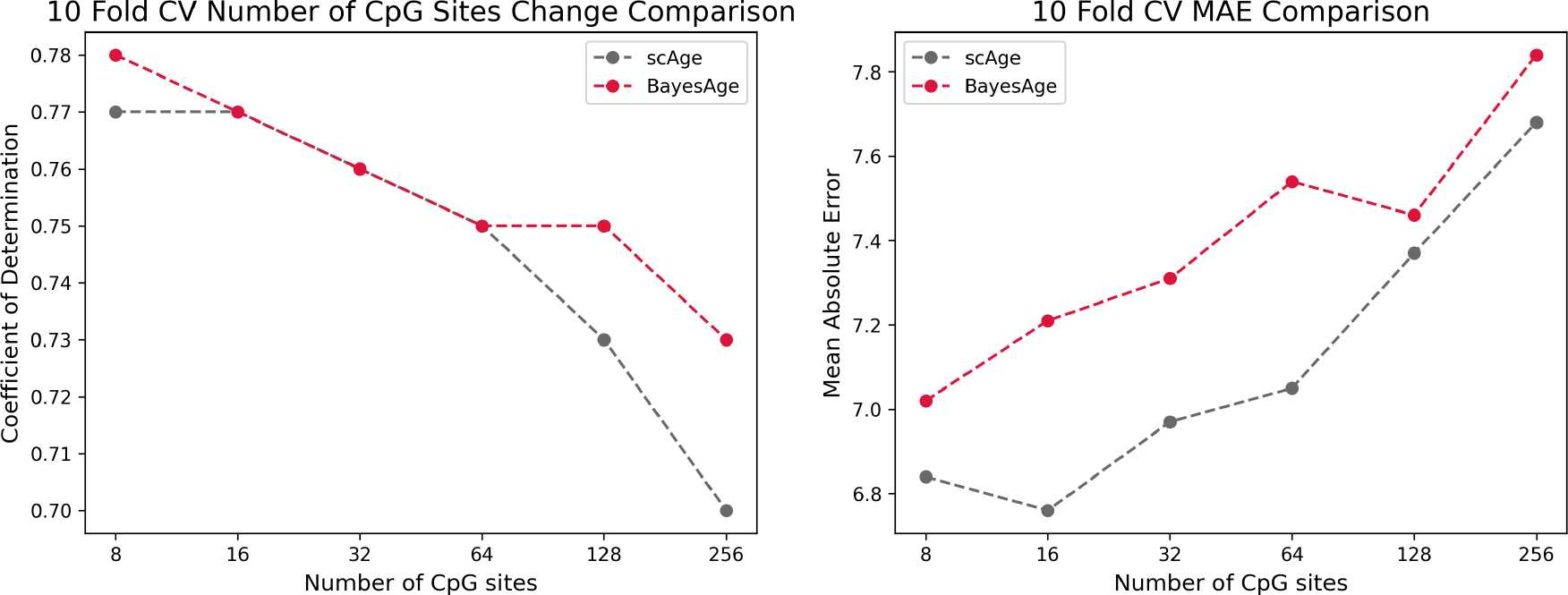
Comparison of the r-squared and MAE of scAge vs BayesAge on a selected number of CpG sites.

While accurately predicting the age of an individual is an important component of these models, another important metric is whether the errors in the prediction show an age bias. In human studies epigenetic clocks are often used to measure the difference between epigenetic and actual age, and many studies have shown that these differences are associated with disease and longevity. However, in order for these age differences to be interpretable, it is important that they do not demonstrate am age dependence. In other words, it is useful for models to generate residuals that are uncorrelated with age.

For this reason we evaluated the residuals of both scAge and BayesAge to identify any age-related biases. Figure 5 shows that the LOWESS fit of scAge, with a tau setting of 0.9, has a nonlinear residual pattern as age varies. By contrast, the BayesAge model was devoid of such age-associated biases in its residuals. The absence of discernible age associated residual patterns is another advantage of of the non-lienar BayesAge approach over scAge.

**Figure 5.**
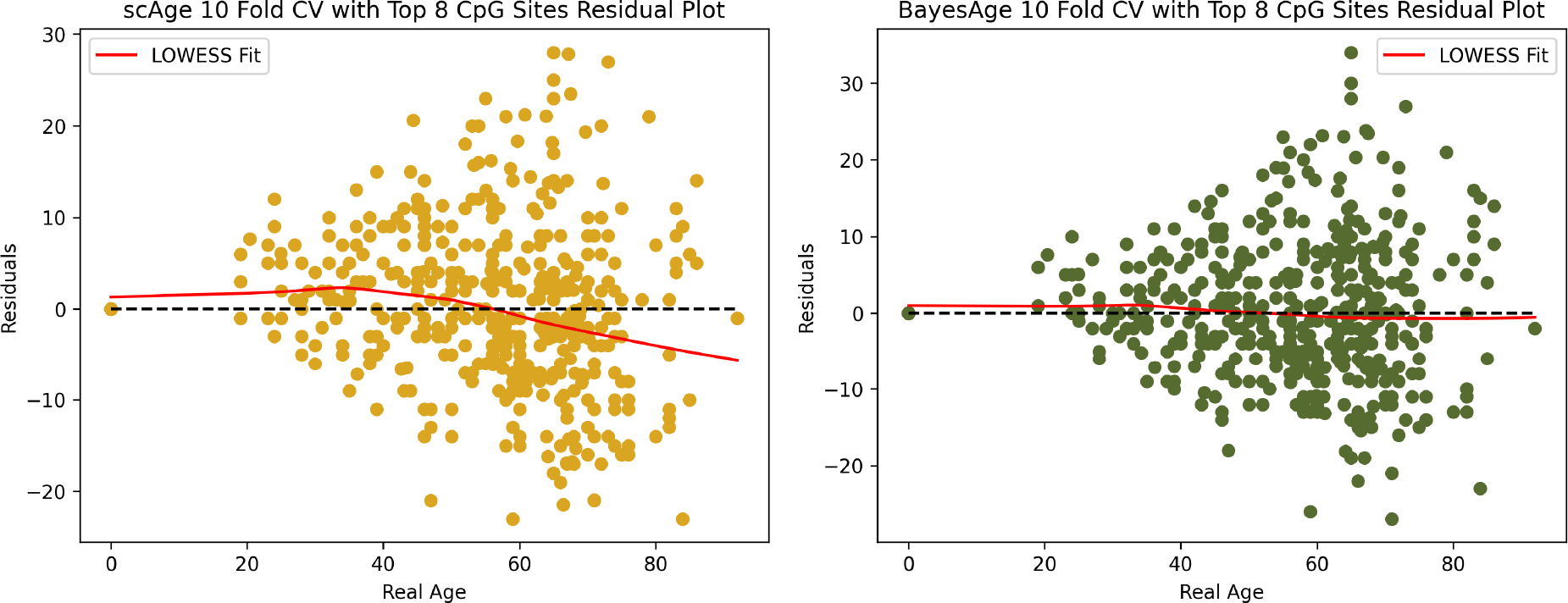
Residual plot analysis of scAge compared to BayesAge.

**Figure 6.**
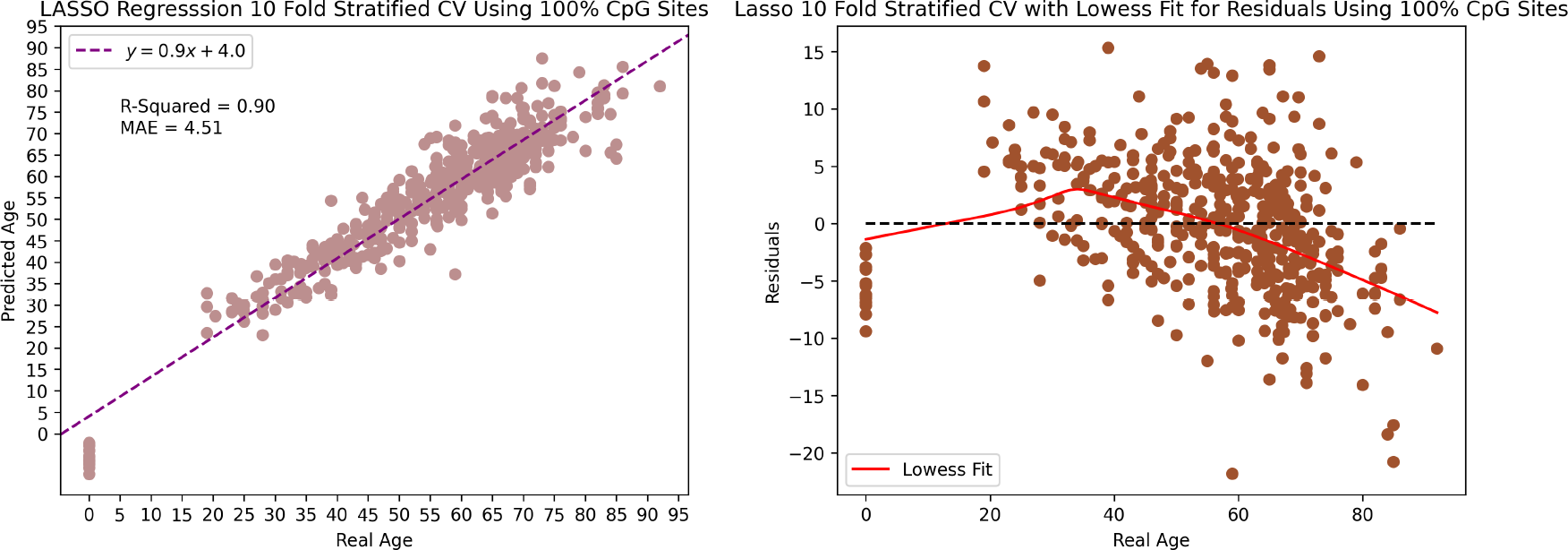
The scatter plot and residual plot of the LASSO model.

**Figure 7.**
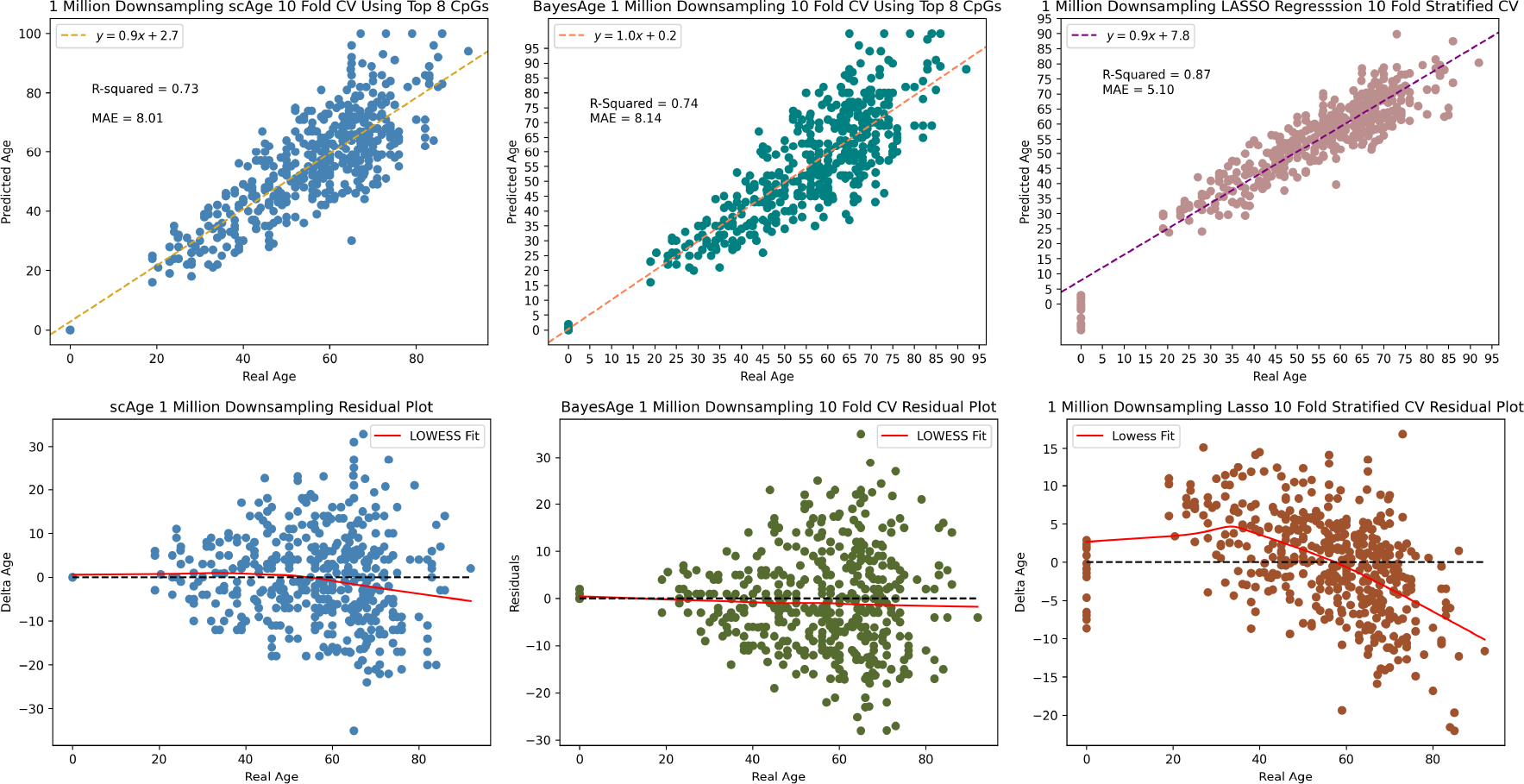
1 million downsampling comparison of scAge vs BayesAge vs LASSO regression. The lowess fit has a tau value of 0.9.

**Figure 8.**
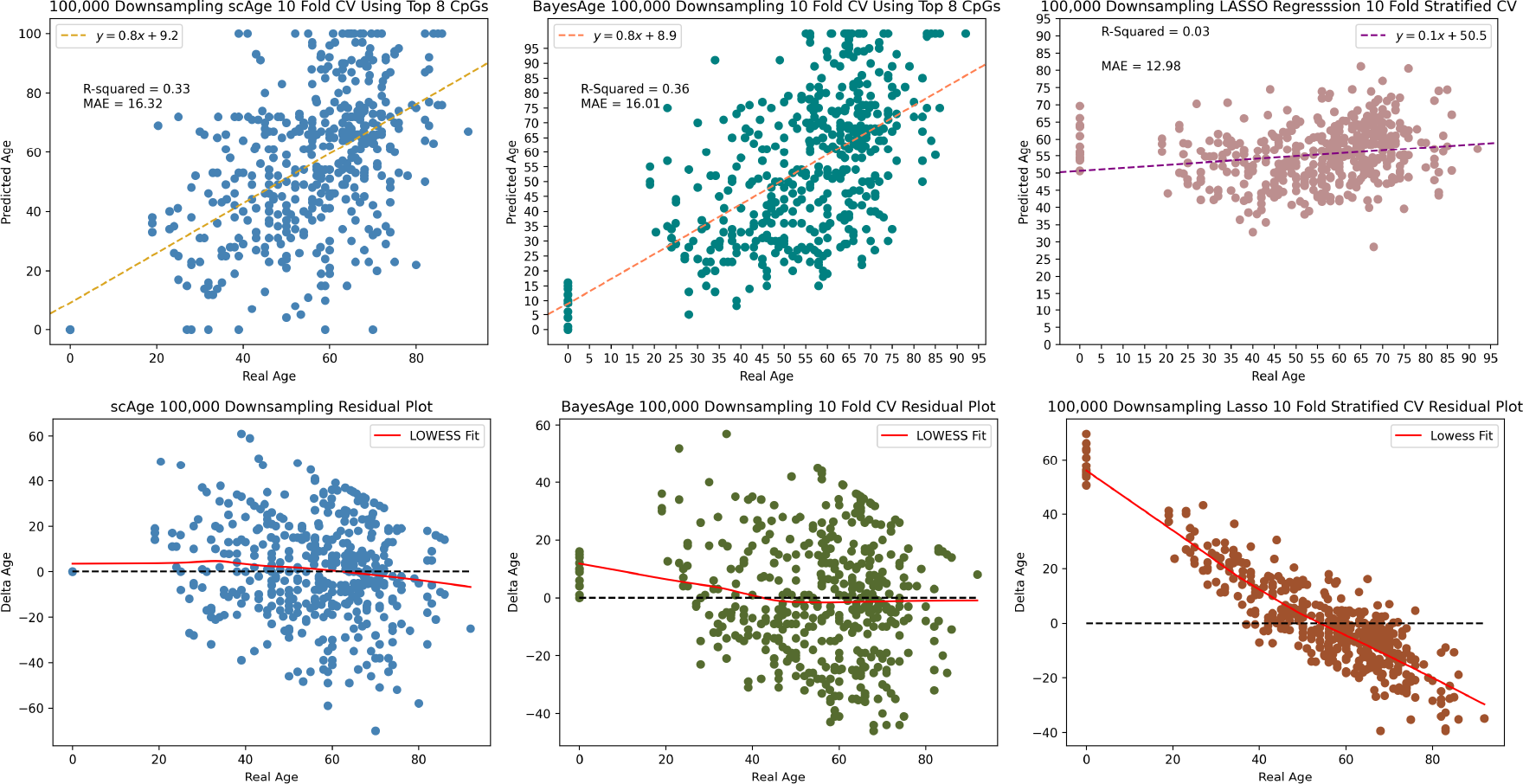
100,000 downsampling comparison of scAge, BayesAge and LASSO regression. LASSO shows a higher bias compared to the MLE models.

Most of the previously published epigenetic clocks have been constructed using regularized regression which constrains the coefficient estimates to zero. Therefore, for comparative purposes, we implemented a LASSO (Least Absolute Shrinkage and Selection Operator) regression trained on our data. Utilizing repeated 10-fold cross-validation, LASSO attained an *r*^2^ value of 90% and a MAE of 4.51 when including all the 46,519 CpG sites in our dataset. Thus the age estimation of the LASSO model outperforms both BayesAge and scAge. However, this comes at a cost of creating a significant bias in the age predictions. This bias is evident as the observed residuals display an age-associated trend that was more pronounced than in scAge. Thus while the age predictions of the LASSO model may be accurate, the residuals, or differences between actual and predicted age, show very strong biases that makes them difficult to interpret.

We used one final metric to evaluate the performance of these three methods, and that involves the measurements of the robustness of the predictions as data is down-sampled. This is important, as bisulfite sequencing data-sets often have varying degrees of coverage. To evaluate model performance at reduced coverage levels, the original bisulfite sequencing data was subjected to random downsampling. At 1 million CpG sites, BayesAge’s *R*^2^ was marginally better at 74%, compared to scAge’s 73%. Despite this, scAge recorded a slightly superior MAE of 8.01, while BayesAge returned 8.14. Notably, the LASSO model outperformed both in terms of *r*^2^ and MAE when compared to other MLE methods. However, the age-associated biases in residuals were distinctly evident for LASSO, more so than the other MLE techniques. Among the three methods, BayesAge demonstrated the least age-related biases.

Upon further reduction to 100,000 reads, BayesAge outperformed the other two methods with an *R*^2^ value of 36% and MAE of 16.01, in comparison to scAge’s respective metrics of 33% and 16.32. The LASSO model was the least resilient to this extreme downsampling, recording a considerably diminished *R*^2^ value of 3% and an elevated MAE of 12.98. Analyzing the residuals revealed significant age-related biases in the LASSO model, in stark contrast to the more consistent patterns observed in scAge and BayesAge models. This comparative analysis, even with signficantly reduced methylation data coverage, underscores BayesAge’s capability to maintain accuracy and limit systemic biases. This positions BayesAge as a useful tool for epigenetic age prediction, particularly in cases with limited data coverage.

## 4 DISCUSSION

We introduced BayesAge, a maximum likelihood estimation framework for predicting epigenetic age from DNA methylation data. BayesAge addresses several limitations found in previous epigenetic age estimation methods such as penalized regression and the Epigenetic Pacemaker. It uses a LOWESS smoothing method to model nonlinear trends in methylation patterns with age, avoiding potential biases that can arise from linear assumptions. The BayesAge model is designed for count-based bisulfite sequencing data. By using a binomial distribution, it effectively models methylation probabilities at each CpG site, accounting for variable coverage depths across different sites and individuals. Notably, BayesAge maintains its performance even in the presence of significant data downsampling.

Our application of age prediction methods to a dataset of 458 individuals indicates that BayesAge generates comparably accurate prediction to the scAge MLE method. However, the residuals of BayesAge show limited age-associated biases, suggesting that our model reflects biological differences in aging that are not correlated with age. This is an important property, as most human epigenetic clock studies are focused on age acceleration rather than age prediction. The fact that many models produce age acceleration estimates that are age associated with age, which confounds their interpretation, and makes it difficult to identify factors that impact the rate of epigenetic aging. By contrast, BayesAge residuals are not age associated and therefore may be more useful for identifying moderators of epigenetic aging.

Another important consideration of epigenetic clocks is that they produce a point estimate of an individual’s age, and it is not possible to obtain a confidence interval on that estimate that accounts for the uncertainty of predictions. To overcome this limitation, we have implemented a simulation framework that allows us to model the range of age estimates that can be generated from a single sample. This simulated data suggests that BayesAge’s predictions have an uncertainty using interquartile range of around 12 years.

In our evaluation of three methods for epigenetic age prediction, we focused on measuring the robustness of predictions as data was down-sampled, a critical consideration given the varying coverage levels often seen in bisulfite sequencing data. When reducing data to 1 million CpG sites, BayesAge displayed a slightly higher *R*^2^ at 74% compared to scAge’s 73%, while scAge had a marginally better MAE at 8.01, versus BayesAge’s 8.14. The LASSO model outperformed MLE methods in *R*^2^ and MAE but exhibited noticeable age-associated biases in residuals. In extreme down-sampling to 100,000 sites, BayesAge surpassed scAge with a higher *R*^2^ (36% vs. 33%) and lower MAE (16.01 vs. 16.32), while the LASSO model’s performance deteriorated significantly (3% *R*^2^, MAE 12.98) with prominent age-related biases. Even with reduced data, BayesAge consistently maintained accuracy and minimized biases, positioning BayesAge as a robust tool for epigenetic age prediction, especially in low-coverage scenarios.

The limitations of our binomial distribution age model provide potential avenues for further research in epigenetic aging frameworks. Future studies might consider using beta-binomial distributions to accommodate overdispersion in methylation probabilities. The model can also be expanded to include biological covariates known to impact methylation, such as gender or smoking habits. On the computational front, implementing optimization methods like expectation-maximization algorithms could enhance efficiency for larger epigenome-wide datasets.

In conclusion, BayesAge offers a comprehensive tool for exploring epigenetic aging dynamics. By addressing the challenges of previous models, BayesAge holds promise for enhancing understanding of aging trajectories across populations. This can lead to insights into factors influencing epigenetic aging, with potential applications in various research areas, from forensics to disease studies.

## CODE AVAILIBILITY

The code for this study is available at: https://github.com/lajoycemboning/BayesAge

